# Facilitating mGluR4 activity reverses the long-term deleterious consequences of chronic morphine exposure

**DOI:** 10.1101/2020.06.27.174771

**Authors:** Jerome AJ Becker, Lucie P Pellissier, Yannick Corde, Thibaut Laboute, Audrey Léauté, Jorge Gandía, Julie Le Merrer

**Author notes:** **Corresponding author:** Julie Le Merrer, PhD Inserm UMR-1253 iBrain, Université de Tours, CNRS, Faculté des Sciences et Techniques, Parc de Grandmont, F-37200 Tours, France.

## Abstract

**Background:** Understanding the neurobiological underpinnings of abstinence from drugs of abuse is critical to allow better recovery and ensure relapse prevention in addicted subjects.

**Methods:** By comparing the long-term transcriptional consequences of morphine and cocaine exposure, we identified the metabotropic glutamate receptor subtype 4 (mGluR4) as a promising pharmacological target in morphine abstinence. We evaluated the behavioral and molecular effects of facilitating mGluR4 activity in abstinent mice.

**Results:** Transcriptional regulation of marker genes of medium spiny neurons (MSNs) in the nucleus accumbens (NAc) allowed best discriminating between 4-week morphine and cocaine abstinence. Among these markers, *Grm4*, encoding mGluR4, displayed down-regulated expression in the caudate putamen and NAc of morphine, but not cocaine, abstinent mice. Remarkably, chronic administration of the mGluR4 positive allosteric modulator (PAM) VU0155041 (2.5 and 5 mg/kg) rescued social abilities, normalized stereotypies and anxiety and blunted locomotor sensitization in morphine abstinent mice. This treatment improved social preference but increased stereotypies in cocaine abstinent mice. Finally, the beneficial behavioral effects of VU0155041 treatment in morphine abstinent animals were correlated with restored expression of key MSN and neural activity marker genes in the NAc.

**Conclusions:** This study is the first report of relieving effects of a pharmacological treatment, chronic administration of the mGluR4 PAM VU0155041, on long-term deleterious consequences of morphine exposure. It illustrates the neurobiological differences between opiate and psychostimulant abstinence and points to pharmacological repression of excessive activity of D2-MSNs in the NAc as a promising therapeutic lever in drug addiction.

## INTRODUCTION

Drug addiction is a chronic psychiatric disorder characterized by loss of control over consumption despite negative consequences (1). Addicted subjects wanting to quit drug use face a true challenge, as protracted abstinence leads to negative affect, anxiety disorders and social withdrawal (2–4) that increase the propensity for relapse (5). A better understanding of the long-lasting neurobiological underpinnings of the abstinent state thus appears invaluable to improve relapse prevention (6–8). Although addiction to most drugs of abuse share similar clinical features, a unitary view of addiction is challenged by different drug-seeking behaviors and brain adaptations across drug classes, particularly for narcotics versus psychostimulants (9–12). The neurobiological mechanisms underlying abstinence and vulnerability to relapse are thus also likely to diverge between drugs.

In previous reports, we have evidenced common long-term transcriptional adaptations that develop within the extended amygdala (EA: central amygdala and bed nucleus of the stria terminalis, CeA and BNST, respectively) upon protracted abstinence to morphine, nicotine, Δ9-tetrahydrocannabinol (THC) and alcohol (13, 14) but not cocaine (14). These adaptations mostly involved genes with an enriched expression in medium spiny GABAergic neurons (MSNs), a key brain substrate for drug reward and addiction (15–17), and notably 14 genes belonging to a huntingtin (HTT)–related gene network (13, 14). Consistent with shared molecular alterations, protracted abstinence from the aforementioned four drugs induced similar behavioral deficits, including impaired social abilities, motor stereotypies and exacerbated anxiety (14, 18–20) that were not observed following cocaine abstinence, except for blunted social preference (14). Remarkably, behavioral deficits evidenced in mice abstinent from morphine, nicotine, alcohol or THC were similar to those detected in mice lacking the mu opioid receptor (*Oprm1*^−/−^), a mouse model of autism (21, 22), and, more generally, in most mouse models of this neurodevelopmental disorder (23, 24). Such similarities may reflect common striatal dysfunction and altered reward processing (25, 26), possibly involving compromised opioid function (27–29). Shared neurobiological mechanisms between drug addiction and autism spectrum disorders interestingly argue for the development of novel therapeutic interventions that would benefit to both pathologies, with the caveat that addiction to cocaine/psychostimulants may diverge from narcotic drugs in their responsiveness to such approaches.

In the present study, we extended our comparison of morphine versus cocaine abstinence-induced transcriptional changes to the caudate putamen (CPu) and nucleus accumbens (NAc), two more brain regions where MSNs represent the main cell type, and analyzed separately the transcriptomes of the CeA and BNST. Not only we confirmed that morphine and cocaine abstinence produce contrasting regulations in gene expression, especially genes with MSN enriched expression, but we detected down-regulated levels of *Grm4* (encoding the presynaptic mGlu4 glutamate receptor -mGluR4) transcripts in the CPu and NAc of morphine, and not cocaine, abstinent animals. Decreased *Grm4* expression retained our attention as another common feature between morphine abstinent and *Oprm1*^−/−^ mice (21, 30), as we evidenced in the latter a rescue of sociability, stereotypic behavior and anxiety levels under chronic facilitation of mGluR4 activity (21). Activation of mGluR4 was shown to inhibit GABA release by indirect, D2 dopamine receptor-expressing MSNs (D2-MSNs), with therapeutic benefit in Parkinson disease (31, 32). Interestingly, negative affect in opioid addiction and dependence may involve D2-MSNs rather than direct, D1 dopamine receptor expressing MSNs (D1-MSNs) (33–35). Here we challenged the hypothesis that chronic administration of VU0155041, a positive allosteric modulator (PAM) and partial agonist of mGluR4, can alleviate the deleterious behavioral and molecular consequences of abstinence to morphine, and presumably not to cocaine. This study provides the first evidence that mGluR4 is a promising pharmacological target to treat the long-term behavioral consequences of opioid use disorder (OUD) and further argues for the involvement of distinct neurobiological processes in opiate versus psychostimulant addiction.

## METHODS AND MATERIALS

### Subjects

In all experiments, we used C57BL6/J male mice aged 8-12 weeks (Charles River, Lyon, France). Animals were group housed and maintained on a 12hr light/dark cycle (lights on at 7:00 AM) at controlled temperature (21±1°C); food and water were available *ad libitum,* except otherwise stated. Animals in the same cage received the same treatment. All experimental procedures were conducted in accordance with the European Communities Council Directive 2010/63/EU and approved by the Comité d’Ethique pour l’Expérimentation Animale de l’ICS et de l’IGBMC (Com’Eth, 2012-033) and Comité d’Ethique en Expérimentation animale Val de Loire (C2EA-19).

### Drugs

Chronic morphine and cocaine treatments were performed as described previously (14) (see time line of experiments, Figure S1). Mice were injected twice daily with vehicle (i.p., NaCl 0.9%) or morphine (i.p., escalating doses from 20 mg/kg to 100 mg/kg over 5 days and one injection on day 6 at 100 mg/kg) or cocaine (s.c., 25 mg/kg during 5,5 days). They were left drug-free for 3 weeks before chronic daily treatment with vehicle (NaCl 0.9%) or VU0155041 (Tocris, Bristol, UK; 2.5 or 5 mg/kg, i.p.) started. This treatment was maintained for 18 consecutive days. All compounds were administered in a volume of 10 ml/kg.

### Antibodies

In Western blot experiments, primary antibodies targeting GAPDH (Rabbit mAb, ref. 2118; RRID: AB_561053) (1:2000) from Cell Signaling Technology and synaptophysin (mouse SYP antibody [D-4], ref. sc-17750; RRID: AB_628311) (1:2000) from Santa Cruz Biotechnology were used at the indicated dilutions. They were combined with the following secondary antibodies: goat anti rabbit IRDye®800CW (LI-COR®, USA, 926-3221; RRID: AB_621843) (1:15000) and goat anti mouse IRDye®680RD (LI-COR®, USA, 926-68070; RRID: AB_10956588) (1:15000).

### Behavioral experiments

As described previously (14, 30), social behavior was explored using the direct social interaction and three-chamber tests, stereotyped/perseverative behavior was assessed by scoring spontaneous motor stereotypies and buried marbles in the marble burying test and by analyzing alternation patterns in the Y-maze, and anxiety-like behavior was evaluated in the novelty-suppressed feeding test. Locomotor activity was recorded using video-tracking (Viewpoint, Lyon, France). Behavioral assays started on week 4 after cessation of morphine or cocaine exposure. On testing days, mice received pharmacological treatment 30 min prior to testing. Detailed protocols can be found in Supplement 1.

### Real-time quantitative Polymerase Chain Reaction analysis

Real-time quantitative Polymerase Chain Reaction (qRT-PCR) analysis was performed on brain samples as described previously (14, 21, 36) (see dissection in Figure S2 and supplementary experimental procedures in Supplement 1).

### Western blot experiments

Mouse brains were dissected 45 min after social interaction and brain regions of interest were punched out (21, 37). Tissues were processed and protein expression was measured as described previously (37) (see supplementary experimental procedures in Supplement 1).

### Statistics

Statistical analyses were performed using Statistica 9.0 software (StatSoft, Maisons-Alfort, France). For all comparisons, values of p<0.05 were considered as significant. Statistical significance in behavioral experiments was assessed using one or three-way analysis of variance (drug, stimulus, treatment and time effects) followed by Newman-Keules post-hoc test. Significance of quantitative real-time PCR (qRT-PCR) results was assessed after transformation using a one-sample t-test, as previously described. Unsupervised clustering analysis was performed on transformed qRT-PCR data (14, 21, 36) using complete linkage with correlation distance (Pearson correlation) for drug, treatment and brain region (Cluster 3.0 and Treeview software). A standard principal component analysis (PCA) was performed on qRT-PCR data and behavioral data if available (21, 36).

## RESULTS

### Deregulated expression of MSN marker genes best discriminated between morphine and cocaine abstinence

We first compared the long-term consequences of a history of morphine or cocaine exposure at transcriptional level across the CPu, NAc, BNST and CeA, by assessing the expression of a set of 76 candidate genes, including the 14 HTT-related genes regulated by morphine abstinence (*Adora2*, *Arpp21*, *Bcl11b*, *Gcnt2*, *Cnr1*, *Drd1a*, *Fam107a*, *Foxp1*,*Gpr88*, *Hpca*, *Nr4a1*, *Pde10a*, *St8sia3, Strip2*)(13), 26 additional marker genes of MSNs (see Table S1) (38–40), as well as key genes involved in GABA, glutamate, monoaminergic and peptidergic neurotransmission and marker genes of neuronal activity and plasticity. We used hierarchical clustering analysis and principal component analysis (PCA) of qRT-PCR data obtained for each drug in each brain region studied (Figure 1, Table S2) to identify groups of genes sharing similar expression profiles.

**Figure 1.**
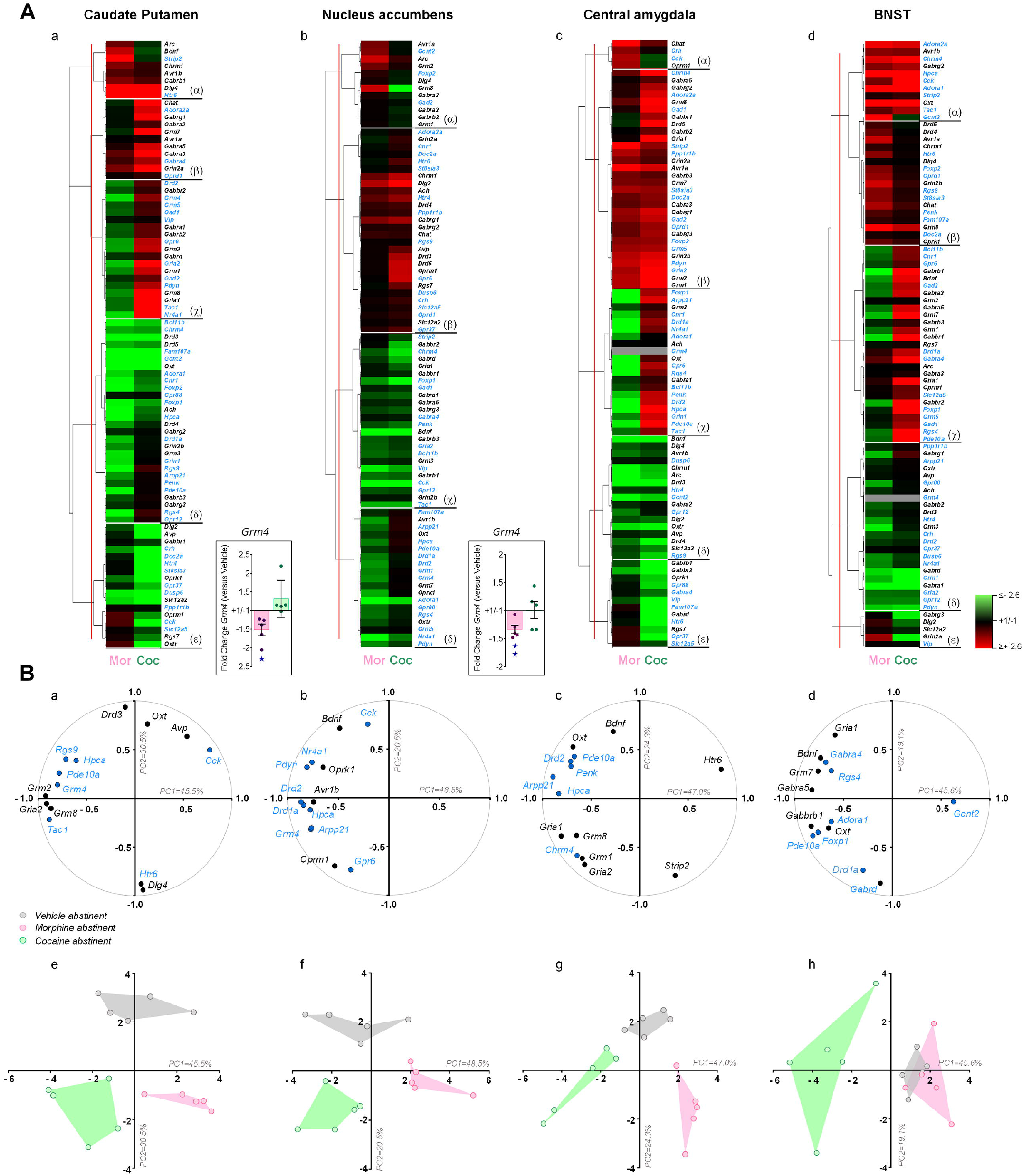
Deregulated expression of MSN marker genes in the CPu, NAc and CeA allows to best discriminate between morphine and cocaine abstinence. **(A)** We used qRT-PCR to compare the expression of 76 candidate genes, among which 14 previously identified HTT-related genes and MSN marker genes (blue characters), in the CPu, NAc, CeA and BNST under morphine and cocaine abstinence conditions compared to their saline counterparts. Hierarchical clustering organized gene expression data in four to five clusters per region. Among these clusters, a limited number showed convergent effects of drugs on transcription (NAc: cluster χ, CeA: cluster χ or BNST: top of cluster α) while most of them illustrated divergent, if not opposed, influence on gene expression, with the CPu displaying the most contrasted morphine versus cocaine effects (clusters β to ɛ). Most HTT-related genes displayed down-regulated expression under morphine condition and gathered in the same clusters in the CPu (cluster δ: 10 genes), NAc (cluster δ: 7 genes) and CeA (cluster δ: 8 genes) together with several other MSN markers (CPu, cluster δ: 7 genes; NAc, cluster δ: 6 genes; CeA, cluster δ: 9 genes). In contrast, the expression of a majority of HTT-related genes was either up-regulated or unchanged under cocaine conditions. In contrast, in the BNST, HTT-related and MSN maker genes were found distributed across all clusters. The expression of *Grm4*, coding for the mGlu4 receptor, was found downregulated in morphine (pink) but not cocaine (green) abstinent animals (framed). It was undetected in the BNST and CeA (grey in clusters). **(B)** A principal component analysis (PCA) performed on qRT-PCR data identified 14 genes (variable’s space) for each region whose expression contributed best to discriminate between morphine (pink) and cocaine (green) abstinence (along PC1, panels a to d). About half of these genes were HTT-related and MSN marker genes in the CPu (panel a), CeA (panel c) and BNST (panel d) and this proportion reached 64% (9 in 14) in the NAc (panel b). PCA analysis successfully segregated morphine, cocaine and vehicle abstinent individuals (subject’s space) in the CPu (panel e), NAc (panel f) and CeA (panel g) but not BNST (panel h), where morphine abstinent subjects partially overlapped with vehicle abstinent animals. Gene expression data are expressed as fold change versus vehicle abstinent mice for each condition (mean ± SEM, n=5 per group). Comparison to vehicle group (*Grm4*): one star p<0.05, two stars p<0.01 (one-sample t test). PC: principal component. Only the two first extracted PCs were considered for representation (PC1 and PC2). qRT-PCR data used for clustering are displayed in Table S2.

In the four regions tested, clustering analysis organized gene expression in four to five clusters (Figure 1A). Among these clusters, a limited number showed convergent effects of these drugs on transcription while most of them illustrated divergent, if not opposed, influence on gene expression, with the CPu displaying the most contrasted morphine versus cocaine effects. Interestingly, most HTT-related genes displayed down-regulated expression under morphine condition and gathered in the same clusters in the CPu, NAc and CeA together with several other MSN markers, while their expression was either up-regulated or unchanged under cocaine conditions. In contrast, in the BNST, these genes were found distributed across all clusters. Thus, the expression of marker genes of MSNs allows to distinguish between morphine versus cocaine abstinence in the CPu, NAc and CeA but not in the BNST.

We then performed a PCA on qRT-PCR data and identified 14 genes for each region whose expression contributed best to segregate experimental conditions (Figure 1B). About half of the identified genes were HTT-related and MSN markers (CPu: panel a, CeA: panel c and BNST: panel d) and this proportion reached 64% (9 in 14) for gene expression in the NAc (panel b), where they tightly correlated with PC1, accounting for most of the variability in the sample. In agreement with clustering results, this analysis successfully segregated morphine, cocaine and vehicle abstinent populations in the CPu (panel e), NAc (panel f) and CeA (panel g) but not BNST (panel h), where morphine abstinent individuals partially overlapped with vehicle abstinent animals. Among the genes whose expression was the most discriminative, we noticed four MSN markers that appeared in at least three regions: *Arpp21, Pde10a, Hpca and Grm4*. *Grm4* retained our attention for showing down-regulated expression under morphine abstinence (and not cocaine abstinence, framed).

### Chronic VU0155041 administration relieved social behavior deficits in morphine and cocaine abstinent mice

We then tested whether facilitating mGluR4 activity by chronically administering a PAM of mGluR4, VU0155041 (2.5 or 5 mg/kg), in morphine abstinent animals would rescue their behavioral deficits. For comparison purpose, we administered the same treatment (VU0155041, 5 mg/kg) to cocaine abstinent mice.

In the direct social interaction test (Figure 2A), morphine abstinent animals displayed a marked deficit in social interaction, as evidenced by significant decrease in the time, number and duration of nose contacts as well as in the number of following episodes, contrasting with increased number of grooming episodes, especially after a social contact. VU0155041 treatment dose dependently (partially at 2.5 mg/kg, completely at 5 mg/kg) normalized these parameters. In contrast, cocaine abstinent animals behaved similarly as vehicle abstinent animals in this test, and VU0155041 treatment had no significant impact on their social behavior, except for an increase in their number of following episodes.

**Figure 2.**
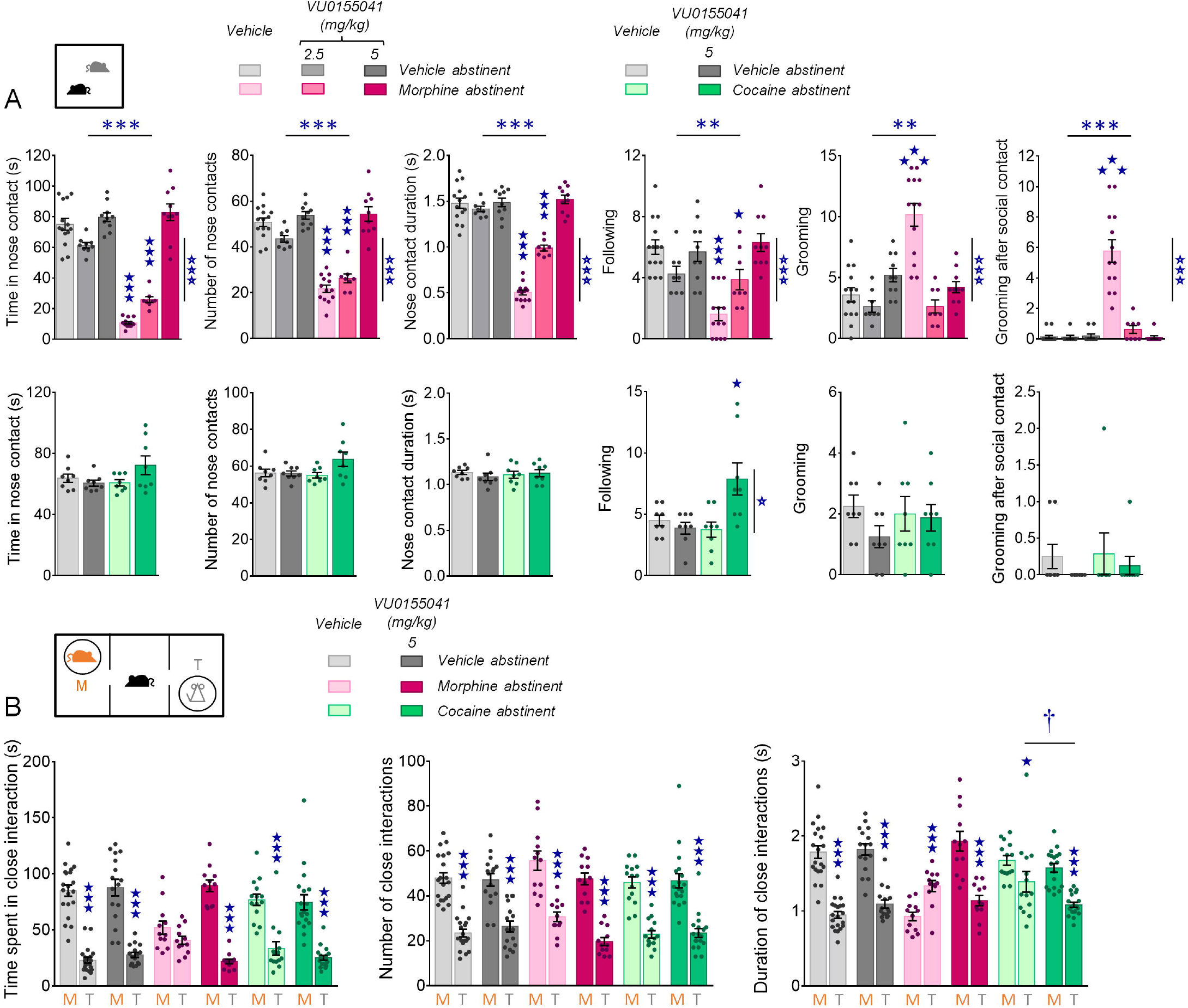
Effects of chronic VU0155041 administration on social behaviors in morphine and cocaine abstinent mice. **(A)** In the direct social interaction test, facilitation of mGluR4 activity in morphine abstinent mice (top row, n=8-14 per group) dose-dependently normalized their time spent in nose contact (abstinence effect: F_1,57_=134.9, p<0.0001; VU0155041 dose effect: F_2,57_=85.9, p<0.0001; abstinence x dose interaction: F_2,57_=56.2, p<0.0001), number of nose contacts (abstinence: F_1,57_=78.6, p<0.0001; dose: F_2,57_=52.0, p<0.0001; abstinence x dose: F_2,57_=28.5, p<0.0001) and duration of nose contacts (abstinence: F_1,57_=148.1, p<0.0001; dose: F_2,57_=71.5, p<0.0001; abstinence x dose: F_2,57_=70.1, p<0.0001), their number of following (abstinence: F_1,57_=9.4, p<0.01; dose: F_2,57_=9.8, p<0.001; abstinence x dose: F_2,57_=13.3, p<0.0001) and grooming episodes (abstinence: F_1,57_=10.8, p<0.01; dose: F_2,57_=19.2, p<0.0001; abstinence x dose: F_2,57_=20.7, p<0.0001), especially these occurring after a social contact (abstinence: F_1,57_=37.1, p<0.0001; dose: F_2,57_=34.6, p<0.0001; abstinence x dose: F_2,57_=35.3, p<0.0001). In cocaine abstinent animals (bottom row, n=8-9 per group), VU0155041 at the dose of 5mg/kg had no significant effect on social behavior, except for an increase in the number of following episodes (abstinence: F_1,28_=4.2, p=0.0502; VU0155041 treatment: F_1,28_=4.9, p<0.05; abstinence x treatment: F_1,28_=8.9, p<0.01). In the three-chamber test, chronic administration of VU0155041 at 5 mg/kg restored social preference in morphine abstinent animals, as shown by rescued longer time spent in close contact (stimulus: F_1,86_= 236.3, p<0.0001; stimulus x abstinence: F_2,86_= 4.1, p<0.05; stimulus x treatment: F_1,86_=9.9, p<0.01; stimulus x abstinence x treatment: F_2,86_= 7.4, p<0.01) and longer-lasting close contacts with the mouse compared to the toy. In cocaine abstinent animals, the difference between close contact duration with the toy and close contact duration with the mouse was significantly increased by mGluR4 PAM treatment (abstinence x treatment: F_2,86_=11.7, p<0.0001; stimulus: F_1,86_= 135.1, p<0.0001; stimulus x abstinence: F_2,86_= 20.3, p<0.0001; stimulus x treatment: F_1,86_=30.4, p<0.0001; stimulus x abstinence x treatment: F_2,86_=23.3, p<0.0001). Results are shown as scatter plots and mean ± sem. Asterisks: abstinence effect, open stars: treatment effect, solid stars: abstinence x treatment interaction, dagger: stimulus x abstinence x treatment, comparison to close contact duration with the toy in vehicle abstinent animals (two-way ANOVA or three-way ANOVA with stimulus as repeated measure, followed by Newman-Keules post-hoc test). One symbol: p<0.05, two symbols: p<0.01, three symbols: p<0.001.

In the three-chamber test (Figure 2B), morphine abstinent animals failed to show preference for interacting with the mouse rather than the toy, as evidenced by similar time spent interacting with both stimuli, and longer duration of their close interactions with the toy than with the mouse. Chronic administration of VU0155041 at 5 mg/kg restored social preference in morphine abstinent animals, rescuing longer time and duration of close contacts with the mouse versus the toy. In cocaine abstinent animals, altered social preference was evidenced by a reduction in the difference between close contact duration with the toy versus the mouse. Chronic VU0155041 significantly shortened the duration of interactions with the toy in these animals. Thus, chronic facilitation of mGluR4 activity relieved social deficits in both morphine and cocaine abstinent animals.

### Chronic VU0155041 treatment normalized stereotypic behavior and anxiety in morphine abstinent mice

We further evaluated the effects of chronic VU0155041 treatment by assessing its influence on other, non-social, long-term behavioral adaptations detected in morphine abstinent animals. We first assessed stereotyped/perseverative behavior in drug abstinent animals. Morphine abstinent mice displayed spontaneous stereotyped grooming, circling and head shakes that were dose-dependently suppressed by chronic VU0155041 administration (Figure 3A). In vehicle and cocaine abstinent mice, however, this treatment increased the frequency of circling episodes. When exploring a Y-maze, morphine abstinent mice displayed impaired spontaneous alternation, mostly due to increased number of perseverative same arm returns. Chronic mGluR4 PAM administration normalized their pattern of exploration; no effect of the treatment was detected in vehicle abstinent animals (Figure 3B). Finally, in the marble-burying test (Figure 3C), chronic VU0155041 reduced the number of marbles buried by morphine abstinent animals to the level of non-treated vehicle abstinent mice. This treatment tended to increase marble burying in vehicle and cocaine abstinent mice. In cocaine abstinent animals, VU0155041 treatment suppressed cocaine-induced increase in spontaneous alternation by restoring the number of alternate arm returns to vehicle values.

**Figure 3.**
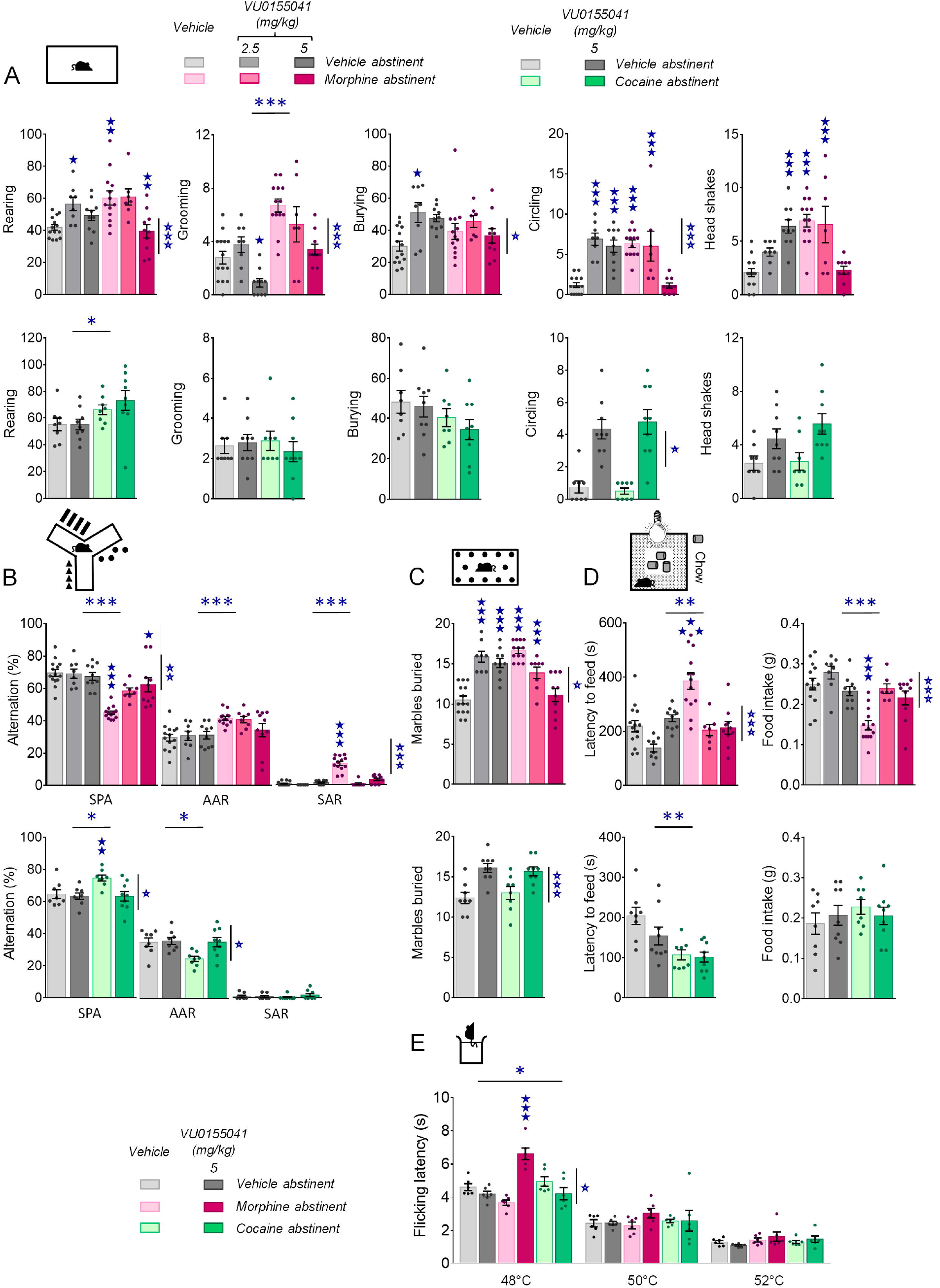
Chronic facilitation of mGluR4 signaling relieved motor stereotypies, perseverative behavior, elevated anxiety and lowered nociceptive thresholds in morphine abstinent mice. **(A)** Under morphine versus vehicle conditions (upper row, n=8-14 per group), chronic VU0155041 administration (2.5 and 5 mg/kg) decreased rearing in morphine-exposed animals (dose: F_2,56_=6.0, p<0.01; abstinence x dose: F_2,56_=7.8, p<0.01), dose-dependently reduced grooming in both morphine and vehicle groups (abstinence: F_1,56_=242.1, p<0.0001; dose: F_2,56_=12.3, p<0.0001), increased burying behavior in vehicle abstinent mice (dose: F_2,56_=4.9, p<0.05; abstinence x dose: F_2,56_=3.3, p<0.05) and finally suppressed circling and head shakes in morphine abstinent mice while increasing the occurrence of these behaviors in vehicle controls (*Circling* -dose: F_2,56_=8.6, p<0.001; abstinence x dose: F_2,56_=29.1, p<0.0001; abstinence x dose: F_2,56_=29.1, p<0.0001; Head shakes - abstinence x dose: F_2,56_=25.9, p<0.0001). Under cocaine versus vehicle conditions (lower row, n=8-9 per group), chronic VU0155041 treatment (5 mg/kg) had no effect on cocaine abstinence-induced increase in rearing frequency (abstinence: F_1,30_=7.4, p<0.05) and increased the frequency of circling episodes (treatment: F_1,30_=7.2, p<0.05). **(B)** In the Y-maze test, chronic VU0155041 administration in morphine abstinent mice (upper panel) dose-dependently restored spontaneous alternation (abstinence: F_1,57_=39.6, p<0.0001; dose: F_2,57_=5.7, p<0.001; abstinence x dose: F_2,57_=8.6, p<0.001) by suppressing same arm returns (abstinence: F_1,57_=67.7, p<0.0001; dose: F_2,57_=41.6, p<0.0001; abstinence x dose: F_2,57_=39.5, p<0.0001). Under cocaine versus vehicle conditions (lower panel), mGluR4 PAM suppressed cocaine abstinence-induced increase in spontaneous alternation (abstinence: F_1,30_=4.3, p<0.05; treatment: F_1,30_=6.9, p<0.05; abstinence x dose: F_1,30_=4.3, p<0.05) by normalizing the number of alternate arm returns in cocaine abstinent mice (abstinence: F_1,30_=5.5, p<0.05; treatment: F_1,30_=5.7, p<0.05; abstinence x dose: F_1,30_=4.2, p=0.05). **(C)** In the marble-burying test, chronic VU0155041 dose-dependently reduced the number of marbles buried by morphine abstinent animals to the level of non-treated vehicle abstinent mice, but increased this number when administered to vehicle abstinent animals (dose: F_2,57_=4.2, p<0.05; abstinence x dose: F_2,57_=7.8, p<0.01). Similarly, VU0155041 increased marble burying in cocaine and vehicle abstinent mice (treatment: F_1,30_=86.8, p<0.0001). **(D)** In the novelty-suppressed feeding test, chronic mGluR4 PAM administration normalized latency to feed and food intake in morphine abstinent animals since the lower dose (abstinence: F_1,56_=11.3, p<0.01; dose: F_2,56_=15.4, p<0.0001; abstinence x dose: F_2,57_=10.5, p<0.001), with no effect in vehicle or cocaine abstinent animals. In the latter, latency to feed was reduced in cocaine abstinent animals (abstinence: F_1,30_=17.5, p=0.001). **(E)** Finally, in the tail immersion test, morphine abstinence tended to decrease nociceptive thresholds measured at 48°C; VU0155041 demonstrated analgesic effects in these animals and only in these animals (abstinence: F_2,30_=4.1, p<0.05; treatment: F_1,30_=7.3, p <0.05; abstinence x treatment: F_2,30_=28.5, p<0.0001). No effect of abstinence nor mGluR4 PAM treatment were detected at 50 and 52°C. Asterisks: abstinence effect; open stars: treatment effect; solid stars: abstinence x treatment interaction, (two-way ANOVA followed by Newman-Keules post-hoc test). One symbol: p<0.05, two symbols: p<0.01, three symbols: p<0.001. AAR: alternate arm returns; SAR: same arm returns; SPA: spontaneous alternation.

We assessed anxiety in abstinent mice using the novelty-suppressed feeding test (Figure 3D). VU0155041 administered at 2.5 or 5 mg/kg normalized the latency to eat in the arena and food intake of morphine-abstinent mice when back in the home cage, with no significant effect in vehicle and cocaine abstinent animals. Finally, we evaluated nociceptive thresholds in morphine and cocaine abstinent mice under vehicle or VU0155041 treatment (5 mg/kg). Morphine abstinent animals showed a tendency for a decrease in their nociceptive thresholds at 48°C, a temperature at which chronic mGluR4 PAM demonstrated analgesic effects in morphine-exposed animals only (Figure 3E).

In conclusion, chronic mGluR4 PAM administration relieved motor stereotypies, perseverative behavior and excessive anxiety and produced analgesia in morphine abstinent mice, with little or no effect in vehicle or cocaine abstinent animals.

### Chronic mGluR4 facilitation inhibited behavioral sensitization to morphine exposure

Then, we assessed morphine and cocaine behavioral sensitization in mice made abstinent from morphine or cocaine and chronically treated with VU0155041 (5 mg/kg) or vehicle. In morphine abstinent mice treated with vehicle, we observed that a single injection of morphine (10 mg/kg) four weeks after cessation of chronic morphine exposure induced a greater locomotor response than in vehicle abstinent animals, demonstrating behavioral sensitization. Chronic VU0155041 administration significantly blunted, although not completely suppressed, this conditioned response (Figure 4A). In cocaine abstinent mice, an acute injection of cocaine (15 mg/kg) induced detectable locomotor sensitization over the first 60 min of the test; chronic facilitation of mGluR4 activity delayed the pic of cocaine-induced locomotor activity (Figure 4B). In conclusion, VU0155041 administration was able to dampen morphine-induced locomotor sensitization in morphine abstinent mice.

**Figure 4.**
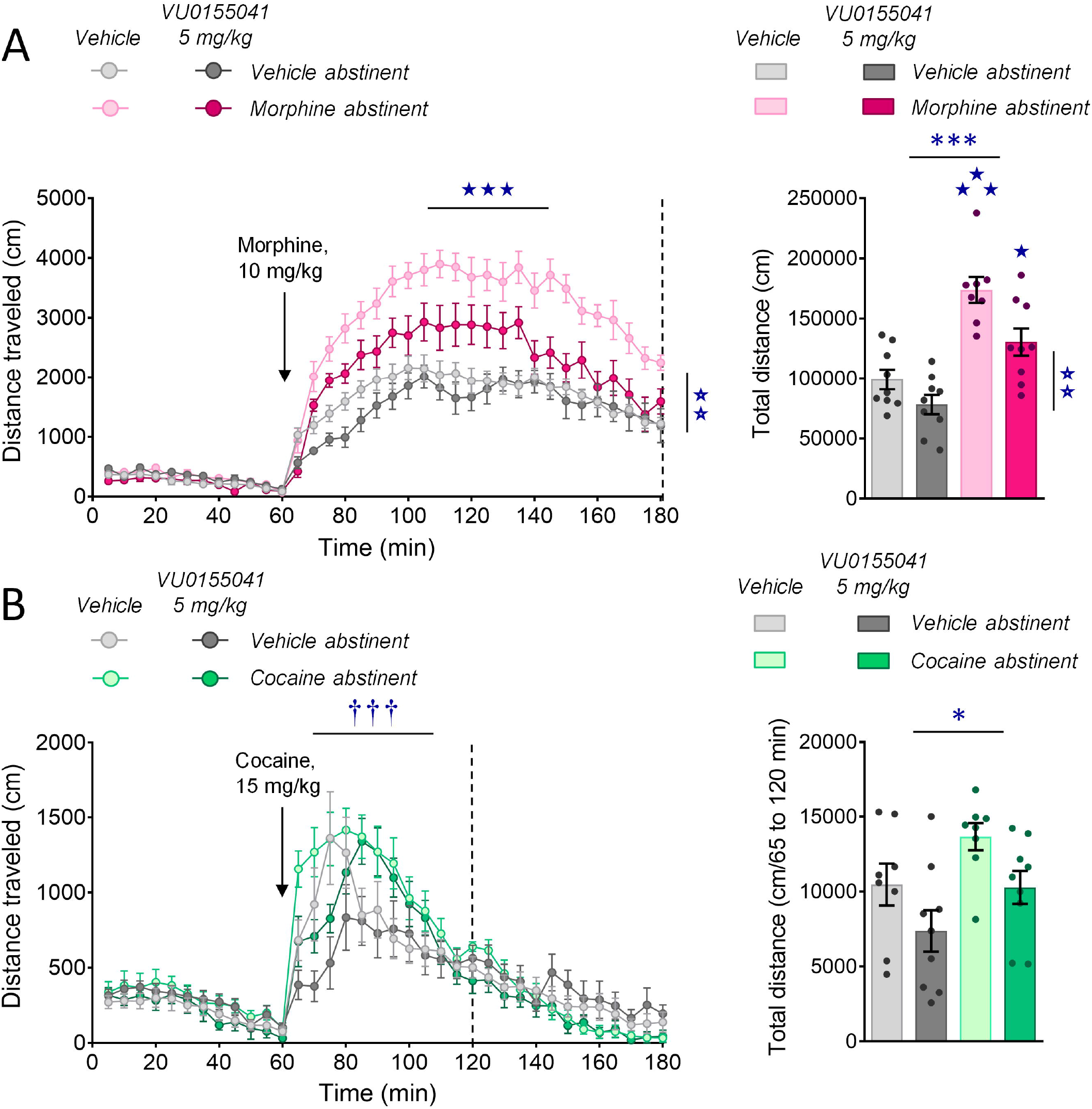
Chronic VU0155041 treatment inhibits morphine-induced locomotor sensitization. **(A)** Morphine abstinent mice displayed locomotor response to acute morphine (10 mg/kg) than vehicle abstinent animals; such behavioral sensitization was significantly decrease upon chronic VU0155041 (5 mg/kg) administration (left panel, n=8-9 per group) as shown by diminished total distance travelled after morphine injection compared to vehicle-treated animals (right panel) (65-180 min; abstinence: F_1,31_=48.5, p<0.0001; treatment: F_1,31_=12.4, p<0.01; abstinence x treatment: F_1,31_=4.8, p<0.05; time: F_23,713_=12.4, p<0.0001; time x abstinence: F_23,713_=4.4, p<0.0001). **(B)** In cocaine abstinent mice, acute cocaine (15 mg/kg) induced detectable locomotor sensitization during the first 60 min of the test; chronic facilitation of mGluR4 activity (VU0155041, 5 mg/kg) delayed the pic of locomotor activity (left panel; n=8-9 per group) but did not significantly decreased cocaine-induced locomotion (right panel) (65-120 min; abstinence: F_1,31_=5.5, p<0.05; time: F_11,352_=18.3, p<0.0001; time x abstinence: F_11,352_=3.0, p<0.001; time x treatment: F_11,352_=5.4, p<0.0001). Results are shown as scatter plots and/or mean ± sem. Asterisk: abstinence effect; open stars: treatment effect; daggers: time x treatment interaction; solid stars: abstinence x treatment interaction (two-or three-way ANOVA followed by Newman-Keules post-hoc test). One symbol: p<0.05, two symbols: p<0.01, three symbols: p<0.001.

### Behavioral improvements in morphine abstinent animals under VU0155041 treatment were correlated with transcriptional changes in the NAc

To gain insight into the molecular mechanisms involved in beneficial effects of VU0155041 treatment, we evaluated its effects on transcriptional regulations induced by morphine abstinence in the CPu, NAC and CeA (where gene expression best characterized morphine versus cocaine abstinence). This experiment was performed following a session of social interaction (Figure S3A), to allow correlations between gene expression and social behavior. We focused on 33 candidate genes, including 16 marker genes of striatal MSNs (belonging or not to the HTT-centered network, blue characters in Figure 5A) as well as genes of the oxytocin/vasopressin system and marker genes of neuronal activity and synaptic activity or plasticity.

**Figure 5.**
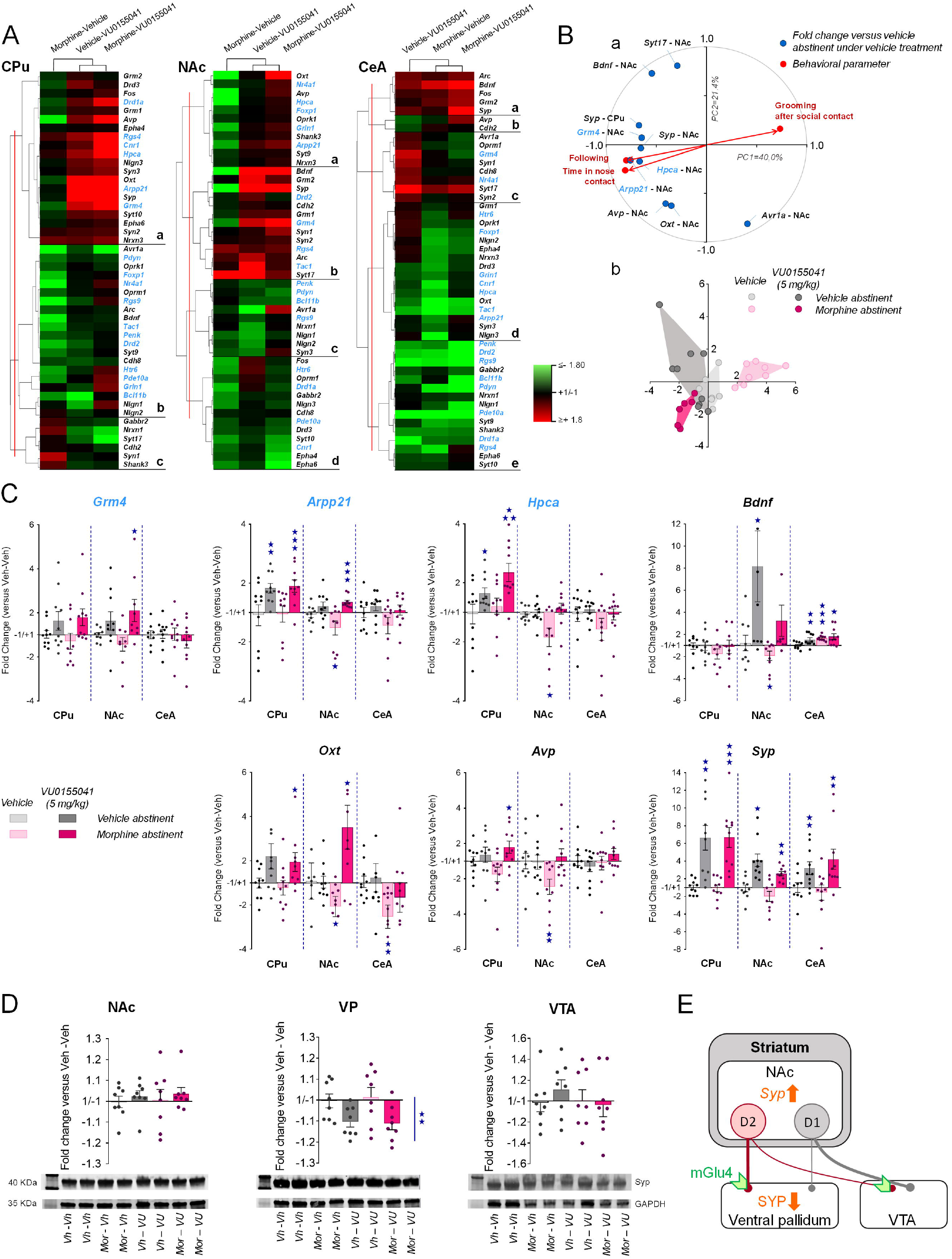
Transcriptional regulations induced in the striatum and amygdala of morphine abstinent animals by chronic facilitation of mGluR4 signaling. We evaluated the effects of chronic VU0155041 treatment (5 mg/kg) on transcriptional changes induced by morphine versus vehicle abstinence in the CPu, NAC and CeA. We focused on 33 candidate genes, among which 16 MSN marker genes (blue characters). **(A)** Clustering analysis organized gene expression in 3 to 5 clusters per region. In the CPu, VU0155041 treatment increased (cluster [a]), normalized (cluster [b]) or decreased (cluster [c]) gene expression as compared to vehicle-morphine condition. In the NAc, animals under VU0155041-morphine condition showed restored (cluster [a] and top of [d]) or increased (cluster [b]) mRNA levels compared to vehicle-morphine mice; no change was detected in cluster [c]. In the CeA, differences between VU0155041-morphine and vehicle-morphine conditions were less contrasted, and mostly observed in cluster [d], where mGluR4 facilitation normalized the expression of most genes. **(B)** PCA analysis identified 10 genes whose expression profile accounted best for the variability in our sample when associated to behavioral parameters (variables, panel a) and that allowed complete segregation between saline-morphine animals and the other experimental groups (subjects, panel b). 9 of these profiles were found in the NAc (*Arpp21, Avr1a, Bdnf, Epha6*, *Grm4*, *Hpca*, *Oxt, Syp, Syt17*) and one in the CPu (*Syp*). **(C)** Most of the identified genes shared similar expression profiles, with down-regulated expression (*Arpp21, Hpca, Oxt, Avp, Bdnf*) or tendency for decreased expression (*Grm4, Syp*) in the NAc of vehicle-morphine mice that was restored (*Hpca, Avp, Bdnf*) or even increased (*Grm4*, *Arpp21, Oxt, Syp*) upon VU0155041 treatment. **(D)** Western blot analysis failed to detect modifications of synaptophysin (SYP) protein levels in the NAc, CPu and VTA but revealed a significant decrease in SYP immunoreactivity in the VP, **(E)** which represents the main projection area of NAc D2-MSNs. Gene (n=9-10 per group) and protein (n=8 per group) expression data are expressed as fold change versus vehicle - vehicle group for each condition (scatter plots and mean ± SEM). Comparison to vehicle - vehicle group: one star p<0.05, two stars p<0.01, three stars: p<0.05 (one-sample t test). PC: principal component. Only the two first extracted PCs were considered for representation (PC1 and PC2). qRT-PCR data used for clustering are displayed in Table S3.

We first performed hierarchical clustering analysis of qRT-PCR data for each brain region to identify groups of genes sharing similar expression patterns (Figure 5A, Table S3). Overall, transcriptional profiles in morphine abstinent mice treated with VU0155041 (morphine-VU0155041) were more similar to those of VU0155041-treated vehicle abstinent (vehicle-VU0155041) mice than vehicle-treated morphine abstinent (morphine-vehicle) mice in the CPu and NAc. Conversely, in the CeA, differences in gene expression patterns between morphine-VU0155041 and morphine-vehicle conditions were the less contrasted. These results indicate that VU0155041 modulated morphine abstinence-induced modifications of gene expression, more significantly in the CPu and NAc.

Next, we performed a PCA on previous qRT-PCR data to identify transcriptional regulations that correlated best with social interaction parameters measured in abstinent animals (Figure 5B). We identified 10 genes whose expression profile tightly correlated with behavioral parameters. Strikingly, 9 of these profiles were found in the NAc and one in the CPu. *Arpp21, Grm4, Hpca* and *Syp* expression profiles in the NAc best correlated with behavioral markers of social motivation. Most of the above genes shared similar expression profiles (Figures 5C), notably in the NAc, where their transcript levels were down-regulated (*Arpp21, Hpca, Oxt, Avp, Bdnf*) or tended to be (*Grm4, Syp*) in morphine-vehicle mice but were restored (*Hpca, Avp, Bdnf*) or even increased (*Grm4*, *Arpp21, Oxt, Syp*) under morphine-VU0155041 condition. Among these genes, *Syp*, coding for the vesicular protein synaptophysin, was the most sensitive to VU0155041 treatment, which induced a marked up-regulation in the three regions tested. In conclusion, mGluR4 facilitation yielded convergent transcriptional effects on a set of genes, normalizing or up-regulating their expression primarily in the NAc.

Intrigued by *Syp* expression data in the striatum, we assessed synaptophysin (SYP) protein levels using western blotting. We dissected out the NAc, as the main substrate of long-term opiate-induced behavioral deficits, its projection sites: ventral pallidum (VP) and ventral tegmental area (VTA), and the neighboring CPu. At behavioral level, we verified again that chronic VU0155041 rescued morphine abstinence-induced deficit in social interaction (Figure S3B). At molecular level, we failed to detect modifications of SYP protein levels in the NAc, CPu and VTA (Figures 5D, S3B, S4 and S5). In the VP, however, we measured a significant decrease in SYP immunoreactivity. PCA analysis (Figure S3C) showed that SYP expression in the VP was negatively correlated with prosocial parameters in the direct social test (panel a), indicating that individuals with low SYP levels in the VP were more prone to interact with their peers (panel b). Thus beneficial effects of chronic VU0155041 were associated with a decrease in SYP levels in the VP, the main projection site of NAc D2-MSNs (Figure 5E).

## DISCUSSION

### Transcriptional regulation of MSN marker genes in the NAc allows best discriminating between morphine and cocaine abstinence

We previously evidenced persistent and contrasted transcriptional regulations in the EA of 4-week morphine versus cocaine abstinent mice for a set of HTT-related genes (13, 14). In the present study, we extended our investigation to a collection of MSN marker genes, to genes involved in GABA, glutamate, monoaminergic and peptidergic neurotransmission and to marker genes of neuronal activity and plasticity. First, we revealed that gene expression patterns in the CeA were more discriminative than those in the BNST to distinguish morphine from cocaine and vehicle abstinence. Then, we evidenced enduring transcriptional modifications in two more brain regions where MSNs represent the main cell type, CPu and NAc. Overall, modified expression of MSN maker genes allowed best segregating morphine from cocaine exposed animals, indicating that abstinence from both drugs significantly impacted this cell population, but in different, if not opposite, directions. A majority of these MSN marker genes displayed common down-regulated expression under morphine condition; they were indifferently genes with enriched expression in D1-(*Drd1a, Pdyn, Tac1*) or D2-MSNs (*Drd2, Foxp1, Penk, Grm4, Rgs4*) or in both MSN populations (*Arpp21, Hpca, Nr4a1, Pde10a, Rgs9*) (38, 39). Thus, prolonged abstinence from morphine, and not cocaine, inhibited the expression of a set of genes in D1- and D2-MSNs. These data substantiate the notion of differential neurobiological substrates involved in opiate versus psychostimulant addiction (9, 14, 41, 42).

The four MSN-enriched genes *Arpp21*, *Pde10a*, *Hpca* and *Grm4* (38, 43–45) captured our interest for displaying highly discriminative patterns of expression between morphine and cocaine conditions. *Arpp21* codes for Cyclic AMP regulated phosphoprotein 21 (ARPP-21) so called regulator of calmodulin signaling (RCS) that modulates striatal calcium and dopamine signaling (46, 47). When phosphorylated, ARPP-21 amplifies D1 but attenuates D2 dopamine receptor signaling in striatal MSNs (47). *Pde10a* encodes the cyclic nucleotide phosphodiesterase 10A (PDE10A) that regulates cGMP and cAMP signaling cascades and striatal function (48, 49). Inhibition of PDE10A stimulates the activity of MSNs, with D2-MSNs being more sensitive to this effect (50, 51). *Hpca* encodes the neural calcium sensor protein hippocalcin, a main contributor of the slow afterhyperpolarization (sAHP) (52, 53). In morphine abstinent rats, sAHP is attenuated in the NAc MSNs, increasing the excitability of a subset of MSNs (54), possibly D2-MSNs (34). Finally, *Grm4*, preponderantly expressed in D2-MSNs (55), encodes the presynaptic mGlu4 glutamate receptor, negatively coupled to adenylyl cyclase, which decreases excitability and reduces neurotransmitter release at axon terminals (56). Therefore, morphine abstinence, by down-regulating the expression of these 4 genes, likely favored an excessive D2-MSN tone. Altered striatofugal balance, notably in the NAc, represents a key cellular mechanism in drug addiction (17, 57). Recent work has elegantly evidenced, by measuring dopamine-induced cAMP signaling in the NAc, that, along repeated opioid exposure, this balance progressively shifts towards preponderant D2-MSN activity, maintained during abstinence (58), as previously observed in morphine withdrawal (34). Exacerbated D2-MSN activity could drive the negative affect associated to morphine dependence and withdrawal, increasing the risk of relapse (33–35). Here, we hypothesized that pharmacological intervention allowing to dampen D2-MSN activity, as by facilitating mGluR4 activity, should relieve long-term behavioral alterations in morphine abstinent mice.

### Chronic facilitation of mGluR4 activity relieves behavioral deficits and dampens behavioral sensitization in morphine abstinent mice

Morphine abstinent mice display a dramatic alteration of their social abilities (14, 18, 20, 59–61). Chronic administration of the mGluR4 PAM VU0155041 dose-dependently normalized social interaction to vehicle-abstinent levels and restored the preference for making more and longer contacts with the mouse versus the toy in the three-chamber test. Chronic VU0155041 similarly restored social abilities in the *Oprm1*^−/−^ mouse model of autism, together with the associated activation of the reward circuit (21). Social interactions are pleasurable events in humans and rodents and therefore activate the brain reward circuit (27, 62, 63). Intriguingly, exacerbated/prolonged D2-MSN tone, notably in the NAc, as observed in morphine abstinent animals (34, 58), was shown to induce aversion (64–66). Hence, excessive D2-MSN activity in morphine abstinent mice may compromise social interactions by hampering their rewarding properties, and chronic VU0155041 treatment would restore social behavior by normalizing this activity.

Besides social impairments, morphine abstinent mice display motor stereotypies, stereotypic marble burying and perseverative behavior in the Y-maze, as previously reported (14). Here we show that chronic VU0155041administration, at the highest dose, suppressed these behaviors. Similarly, VU0155041 alleviated stereotypies in *Oprm1*^−/−^ mice (21) and relieved stereotypies in mice lacking the *Elfn2* gene (*Elfn2*^−/−^), which also display autistic-like symptoms (67). Interestingly, stereotypic behavior was shown to involve NAc D2-MSNs in another mouse model of autism, Neuroligin-3 knockout mice (68). Hence, VU0155041 may reduce stereotypies in morphine abstinent mice by repressing D2-MSN activity.

We next verified that conflict anxiety was exacerbated in morphine abstinent mice, as evidenced by increased latency to eat in the novelty-suppressed feeding test (14). From the lowest dose we tested, chronic VU0155041 normalized anxiety levels, as previously observed in *Oprm1*^−/−^ and *Elfn2*^−/−^ mice (21, 67). This result is consistent with previous reports of facilitating mGluR4 activity having anxiolytic effects in mice (69–71) and rats (72). In addition, VU0155041 treatment produced analgesia only in morphine abstinent mice whose tendency for lowered nociceptive thresholds is reminiscent of withdrawal symptoms (73). Stimulation of mGluR4 produces analgesia under chronic pain or inflammatory conditions only (74, 75). Interestingly, opioid-induced hyperalgesia was shown to involve an activation of the glutamatergic system (76) that may represent the neurobiological substrate for analgesic effects of VU0155041 in morphine abstinent mice. Finally, morphine-exposed mice displayed significant locomotor sensitization to morphine after prolonged abstinence, consistent with previous findings (14, 77), which was dampened by VU055041. Morphine-induced sensitization involves D1 receptor activation in striatal D1-MSNs (77–79), whose activity may be impacted by mGluR4 either directly in D1-MSNs (80, 81) or through corticostriatal terminals forming synapses at D1-MSNs (82). However, an effect of VU0155041 at D2-MSNs cannot be ruled out, as they also contribute to drug-induced sensitization (83, 84).

Collectively, blunted social abilities, stereotyped behavior, elevated anxiety and low nociceptive thresholds in morphine abstinent mice fit with clinical reports of social withdrawal, impaired cognitive flexibility and high comorbidity with anxiety disorder and depression in patients with a history of chronic opioid exposure (85–92). Importantly, such negative affect, combined with conditioned effects of opioids, are considered as the two major causes of relapse in Opioid Use Disorder (93, 94); they were both significantly alleviated in morphine abstinent mice under VU0155041 treatment.

In contrast with morphine, cocaine abstinence had little impact on sociability, as previously shown (14), and VU0155041 treatment either preserved or even improved these abilities. Chronic VU0155041, however, induced motor stereotypies in these animals as it did in vehicle abstinent mice, possibly by tilting the D1/D2-MSN balance towards D1-MSN activity, which can trigger stereotypic behaviors (95). These results further illustrate the neurobiological differences between opiate and cocaine abstinence.

### VU0155041-induced transcriptional changes in the NAc are coherent with a repression of D2-MSN activity and restauration of social reward

We proposed above that mGluR4 PAM treatment relieved social abilities in morphine abstinent mice by repressing D2-MSN activity in the NAc. Our qPCR results in morphine abstinent mice treated or not with VU0155041 were consistent with this hypothesis, by showing notably that modified expression of a set of genes in the NAc upon VU0155041 treatment correlated best with social interaction parameters. VU0155041 restored the expression of *Grm4*, *Arpp21* and *Hpca*, whose respective encoded protein can dampen D2-MSN activity. Similarly, NAc *Bdnf* expression was down-regulated under prolonged morphine abstinence (96), but restored upon VU0155041 treatment. Reduced BDNF signaling in the NAc would decrease GABAergic activity specifically in D1-MSNs (97), favoring excessive D2 tone. Thus, restoring *Bdnf* expression under mGluR4 PAM treatment may have participated in rebalancing striatofugal activities. Moreover, striatal *Syp* mRNA levels, especially in the NAc, increased under VU0155041 treatment in correlation with social interaction. VU0155041-induced rise in *Syp* expression appears a compensatory mechanism, as SYP protein levels were decreased in the VP, the main projection area of NAc D2-MSNs. The mGlu4 receptor is located presynaptically at axon terminals where its activation represses neurotransmitter release. Thus, reduced expression of SYP in the VP likely resulted from the inhibitory action of mGluR4 PAM treatment on GABA release from indirect MSNs in this region. These data point to repression of D2-MSN activity in the NAc as a key substrate supporting social effects of mGluR4 facilitation in morphine abstinent mice.

The NAc is a hub region for the integration of reward processes (98–100), a key brain substrate in drug addiction (101, 102) and finally a key substrate for social behavior. Consistent with their pleasurable properties, social stimuli activate the NAc in humans (103, 104) and this activation is blunted in patients with ASD (27), depression (105), schizophrenia (106) or OUD (107) who display impaired social abilities. In rodents, activating neuromodulator or neuropeptide receptors in the NAc stimulates or drives social behavior (108–111) and models of ASD, schizophrenia or depression show abnormal volume, connectivity and/or function of the NAc (112–117). In our study, *Fos* expression induced by social exposure was dampened in morphine abstinent mice, suggesting blunted social reward, and rescued upon VU0155041 treatment, as in *Oprm1*^−/−^ mice (21). Moreover, NAc expression of *Oxt* and *Avp* transcripts, plausibly transported from the hypothalamus (118), was restored by mGluR4 PAM treatment. The former gene codes for oxytocin, critically involved in mediating social reward in the NAc (109, 119); the role of vasopressin (encoded by *Avp*) in the NAc remains poorly understood, but vasopressin dysregulation has been highlighted in autism (120). Together, these results argue for a restoration of social reward under VU0155041 treatment, likely through a resetting the D1/D2-MSN balance in the NAc. Further investigations will be required, however, to confirm this hypothesis and also to explore the role of NAc and extra-NAc mGluR4 populations in regulating social behavior. Of note, *Grm4* expression was not down-regulated upon VU0155041 administration, suggesting limited tolerance to pharmacological treatment.

### Conclusion

The present study is the first report that facilitating glutamate mGluR4 activity can relieve the deleterious long-term behavioral and molecular consequences of opiate, but not cocaine, abuse. These results highlight the neurobiological differences between opiate and psychostimulant abstinence, arguing against a unitary view of drug addiction (9, 41). They also substantiate the view of negative affect in opiate abstinence as resulting from excessive activity of striatal D2-MSNs, more specifically in the NAc (33–35, 58). In this context, pharmacological compounds repressing striatopallidal activity appear as promising candidates for the treatment of OUD, possibly by restoring “liking” (121) for non-drug, notably social, stimuli. More broadly, opiate addiction sharing striking phenotypic and neurobiological features with other diseases considered as “reward deficiency syndromes”, such as autism or depression (27, 116, 122, 123), our work opens promising avenues towards the development of common therapeutic strategies for these pathologies.

## Supporting information

Table S2 - qPCR results

Table S3 - qPCR results

Table S1 - list of primers

## Acknowledgments

We are grateful to Dr B.L. Kieffer for fruitful discussions and support at an early stage of this project. We thank G. Duval and D. Memedov for animal care and technical assistance (Institut de Génétique et de Biologie Moléculaire et Cellulaire). We thank the Experimental Unit PAO-1297 (EU0028, Animal Physiology Experimental Facility, DOI: 10.15454/1.5573896321728955E12) from the INRAE-Val de Loire Centre for animal breeding and care. This work was supported by the Institut National de Recherche pour l’Agriculture, l’alimentation et l’Environnement (INRAE), Centre National de la Recherche Scientifique (CNRS), Institut National de la Santé et de la Recherche Médicale (Inserm), and Université de Tours. We thank Région Centre (ARD2020 Biomédicament – GPCRAb) and LabEx MabImprove for financial support. J.L.M. acknowledges postdoctoral fellowship from the Fondation Université de Strasbourg, generously granted by Pierre Fabre Laboratories. L.P.P. acknowledges postdoctoral support from the Marie-Curie/AgreenSkills Program.

## Author contributions

J.A.J.B., J.G., L.P.P. and J.L.M. designed the experiments. J.A.J.B., L.P.P, J.G., T.L.,mA.L. and J.L.M. performed behavioral experiments. J.A.J.B., Y.C. and L.P.P. performed qRT-PCR experiments. Y.C. performed western blot experiments. J.A.J.B., L.P.P., J.G., Y.C. and J.L.M. analyzed the data. J.A.J.B. and J.L.M. wrote the article. All authors discussed the results and commented on the manuscript.

