## Supplementary information for "Facilitating mGluR4 activity reverses the long-term deleterious consequences of chronic morphine exposure"

### Supplementary experimental procedures

#### Behavioral experiments

Experiments in morphine and cocaine abstinent (effects of VU0155041 treatment), were performed in a battery, except for the three-chamber test performed in separate cohorts of naïve mice. Direct social interaction, novel object recognition and novelty suppressed feeding were performed in 4 equal square arenas (50 x 50 cm) separated by 35cm-high opaque grey Plexiglas walls over a white Plexiglas platform (View Point, Lyon, France). Stimulus mice used for the three-chamber test were 8-10-week-old male grouped-housed wild-type mice, socially naive to the experimental animals.

***Social abilities***

**Direct social interaction test.** On testing day, a pair of unfamiliar mice (not cage mates, age-, sex- and treatment-matched) was introduced in each arena for 10 min (15 lx). Each arena received a black plastic floor (transparent to infrared) covered with lightly sprayed fresh sawdust to limit anxiety. The total amount of time spent in nose contact (nose-to-nose, nose-to-body or nose-to-anogenital region), the number and duration of these contacts, grooming episodes (allogrooming), notably ones occurring immediately (<5s) after a social contact, as well as the number of following episodes were scored *a posteriori* on video recordings (infrared light-sensitive video camera) (1-3) using an ethological keyboard (Labwatcher®, View Point, Lyon, France) by trained experimenters and individually for each animal. The mean duration of nose contacts was calculated from previous data (4-6).

**Three-chamber social preference test.** The test apparatus consisted of a transparent acrylic box (exterior walls blinded with black plastic film); partitions divided the box into three equal chambers (40 x 20 x 22.5 cm). Two sliding doors (8 x 5 cm) allowed transitions between chambers. Cylindrical wire cages (18 x 9 cm, 0.5 cm diameter-rods spaced 1 cm apart) were used to contain the mouse interactor and object (soft-toy mouse). The test was performed in low-light conditions (15 lx) to minor anxiety. Stimulus wild-type mice were habituated to confinement in wire cages for 2 days before the test (20 min/day). On testing day, the experimental animal was introduced to the middle chamber and allowed to explore the whole apparatus for a 10-min habituation phase (wire cages empty) after the sliding doors were raised. The experimental mouse was then confined back in the middle-chamber while the experimenter introduced an unfamiliar wild type age and sex-matched animal into a wire cage in one of the side-chambers and a soft toy mouse (8 x 10 cm) in the second wire cage as a control for novelty. Then the experimental mouse was allowed to explore the apparatus for a 10-min interaction phase. The time spent in each chamber, the time spent in close contact (nose or paw contact) with each wire cage (empty: habituation; containing a mouse or a toy: interaction), as well as the number of these close contacts were scored *a posteriori* on video recordings using an ethological keyboard (Labwatcher®, View Point, Lyon, France) by trained experimenters. The mean duration of close contacts was calculated from these data (4-6). The relative position of stimulus mice (versus toy) was counterbalanced between groups (1, 7).

***Stereotyped behaviors***

**Motor stereotypies.** To detect motor stereotypies in mutant versus wild-type animals, mice were individually placed in clear standard home cages (21×11×17 cm) filled with 3-cm deep fresh sawdust for 10 min (8). Light intensity was set at 30 lux. Trained experimenters scored numbers of head shakes, as well as rearing, burying, grooming, circling episodes and total time spent burying by direct observation.

**Y-maze exploration.** Spontaneous alternation behavior was used to assess perseverative behavior (9-11). Each Y-maze consisted of three connected Plexiglas arms (15x15x17 cm) covered with distinct wall patterns (15 lx). Floors were covered with lightly sprayed fresh sawdust to limit anxiety. Each mouse was placed at the center of a maze and allowed to freely explore this environment for 6 min. The pattern of entries into each arm was quoted on video-recordings. Spontaneous alternations (SPA), i.e. successive entries into each arm forming overlapping triplet sets, alternate arm returns (AAR) and same arm returns (SAR) were scored, and the percentage of SPA, AAR and SAR was calculated as following: total / (total arm entries -2) * 100 (1, 12).

**Marble-burying.** Marble burying was used as a measure of perseverative behavior (13). Mice were introduced individually in transparent cages (21×11×17 cm) containing 20 glass marbles (diameter: 1.5 cm) evenly spaced on 4-cm deep fresh sawdust. To prevent escapes, each cage was covered with a filtering lid. Light intensity in the room was set at 40 lux. The animals were removed from the cages after 15 min, and the number of marbles buried more than half in sawdust was quoted (1, 12).

***Anxiety-like behavior***

**Novelty-suppressed feeding.** Novelty-suppressed feeding (NSF) was measured in 24-hr food-deprived mice, isolated in a standard housing cage for 30 min before individual testing. Three pellets of ordinary lab chow were placed on a white tissue in the center of each arena, lit at 60 lx. Each mouse was placed in a corner of an arena and allowed to explore for a maximum of 15 min. Latency to feed was measured as the time necessary to bite a food pellet. Immediately after an eating event, the mouse was transferred back to home cage (free from cage-mates) and allowed to feed on lab chow for 5 min. Food consumption in the home cage was measured (1, 12, 14).

***Locomotor activity under pharmacological challenge***

Locomotor activity was assessed in clear Plexiglas boxes (21 × 11× 17 cm) placed over a white Plexiglas infrared-lit platform. Light intensity of the room was set at 15 lx. The trajectories of the mice were analyzed and recorded via an automated tracking system equipped with an infrared-sensitive camera (Videotrack; View Point, Lyon, France). To focus on forward activity, only movements which speed was over 6 cm/s were taken into account for the measure of locomotor activity.

Behavioral testing started when the animals were placed in the activity boxes for a 60 min-habituation period. Habituation allowed the animals to reach a low and stable level of basal activity and ensured reliable subsequent measure of drug-induced locomotor effects. Mice were then injected with either vehicle (NaCl 0.9%) morphine

(10 mg/kg, s.c.) or cocaine (25 mg/kg, s.c.), and locomotor activity was monitored for further 120 min.

#### Real-time quantitative PCR analysis

*Experiment 1.* Separate cohorts of morphine versus vehicle and cocaine versus vehicle abstinent animals were prepared for qRT-PCR experiments (Figure S1). Brains were removed and placed into a brain matrix (ASI Instruments, Warren, MI, USA). CPu, NAc, BNST and CeA were punched bilaterally out from 1mm-thick slices. Tissues were immediately frozen on dry ice and kept at -80°C until use. For each structure of interest and treatment condition, samples were prepared from five animals processed individually (n=5).

*Experiment 5*. A cohort of morphine versus vehicle abstinent mice was submitted to a direct social interaction test and sacrificed 45 min after the beginning of behavioral assay for qRT-PCR experiments (Figure S1). Brains were removed and placed into a brain matrix (ASI Instruments, Warren, MI, USA). CPu, NAc, and CeA were punched bilaterally out from 1mm-thick slices. Tissues were immediately frozen on dry ice and kept at -80°C until use. For each structure of interest and treatment condition, samples were prepared from eight animals processed individually (n=8).

RNA was extracted and purified using the MIRNeasy mini-kit (Qiagen, Courtaboeuf, France). cDNA was synthetized using the first-strand Superscript II kit (Invitrogen®, Life Technologies, Saint Thomas, France) (15). qRT-PCR was performed in quadruplets on a LightCycler 480 Real-Time PCR (Experiment 1: Roche, Manheim, Germany; Experiment 2: CFX384 Biorad, Marnes-la-Coquette, France) using iQ-SYBR Green supermix (Bio-Rad, Marnes-la-Coquette, France) kit with 0.25µl cDNA in a 12.5µl final volume. Gene-specific primers were designed using Primer3 software to obtain a 100- to 150-bp product (Table S1). Relative expression ratios were normalized to the level of actin and the $2^{-\Delta\Delta Ct}$ method was applied to evaluate differential expression level.

#### Western blot experiments

*Experiment 4*. Mouse brains were dissected 45 min after social interaction (Figure S1) and placed into a brain matrix (ASI Instruments, Warren, MI, USA). Caudate putamen (CPu), nucleus accumbens (NAc), ventral pallidum (VP) and ventral tegmental area (VTA) were punched out. Tissues were immediately homogenized on RIPA buffer (Cell Signaling Technology) containing 1mM of PMSF and 1X Protease/Phosphatase Inhibitor Cocktail, incubated 30 min on ice, centrifuged at 10000 g for 10 min at 4°C, and supernatants were prepared for Western blotting by adding the appropriate volume of 4X Laemmli sample buffer (NuPAGETM LDS Sample buffer NP0007, ThermoFisher Scientific) containing 10% β-mercaptoethanol. The samples were then frozen at -20°C.

For western blot analysis, samples were denaturated at 95°C for 15 min and cleared by centrifugation at 11,000 g for 5 min. Supernatant was loaded onto SDS-acrylamide 4–15% Mini-PROTEAN® TGX™ Precast Protein Gels (BioRad) and proteins were transferred to nitrocellulose membranes with Trans-Blot Turbo™ RTA transfer kit (BioRad) using the Trans-Blot Turbo™ Transfer System (BioRad). Membranes were blocked with 5% (w/v) milk powder diluted in TBS-T (Tris-buffered saline with 1% (v/v) Tween20) for 1h and then incubated at 4ºC overnight with the primary antibody. Blots were finally incubated, in the dark, with fluorescent secondary antibodies, at room temperature. Revelation was performed using the infrared scanner Odyssey® CLx (LI-COR Biotechnology). Quantification was performed using Image Studio software.

### Supplementary Figures

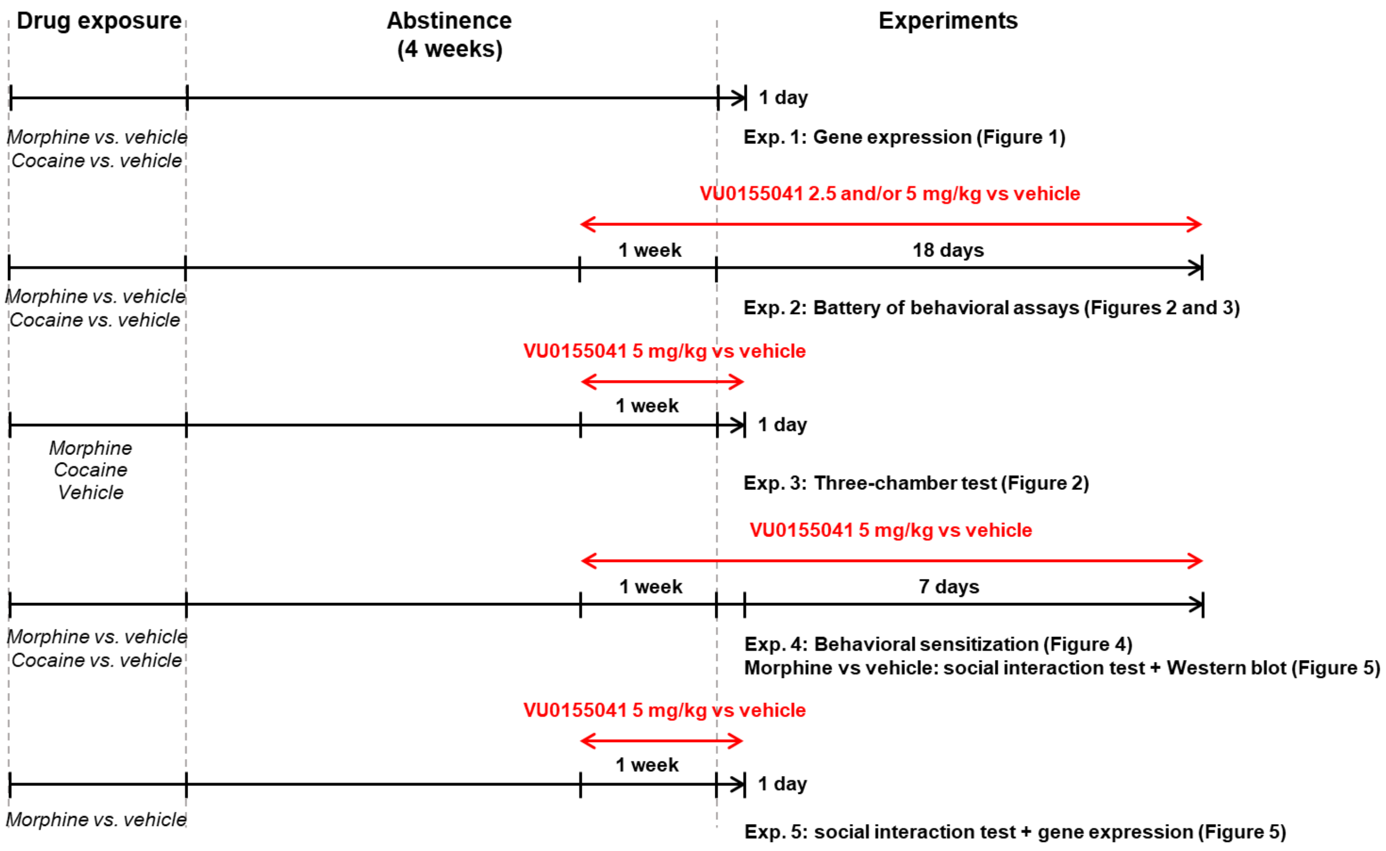


**Figure S1. Time line of experiments.** Exp: experiment.

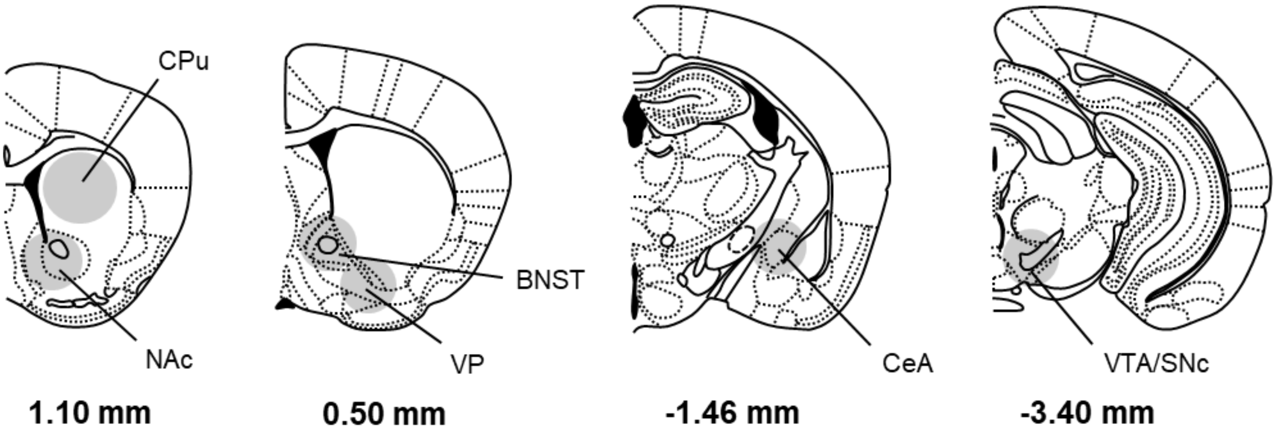


**Figure S2. Schematic representations depict brain regions dissected for gene expression study and western blot experiments.** CPu, NAc, BNST, VP, CeA, and VTA/SNc were punched on 1-mm thick brain slices (CPu: one punch/side, ⌀ 2 mm; NAc, BNST, VP, CeA, and VTA/SNc: one punch/side, ⌀ 1.25 mm). Coordinates refer to bregma. BNST: Bed Nucleus of the Stria Terminalis; CPu: Caudate Putamen;

CeA: Central Amygdala; NAc: Nucleus Accumbens; SNc: Substancia Nigra, pars compacta; VP: ventral pallidum; VTA: Ventral Tegmental Area.

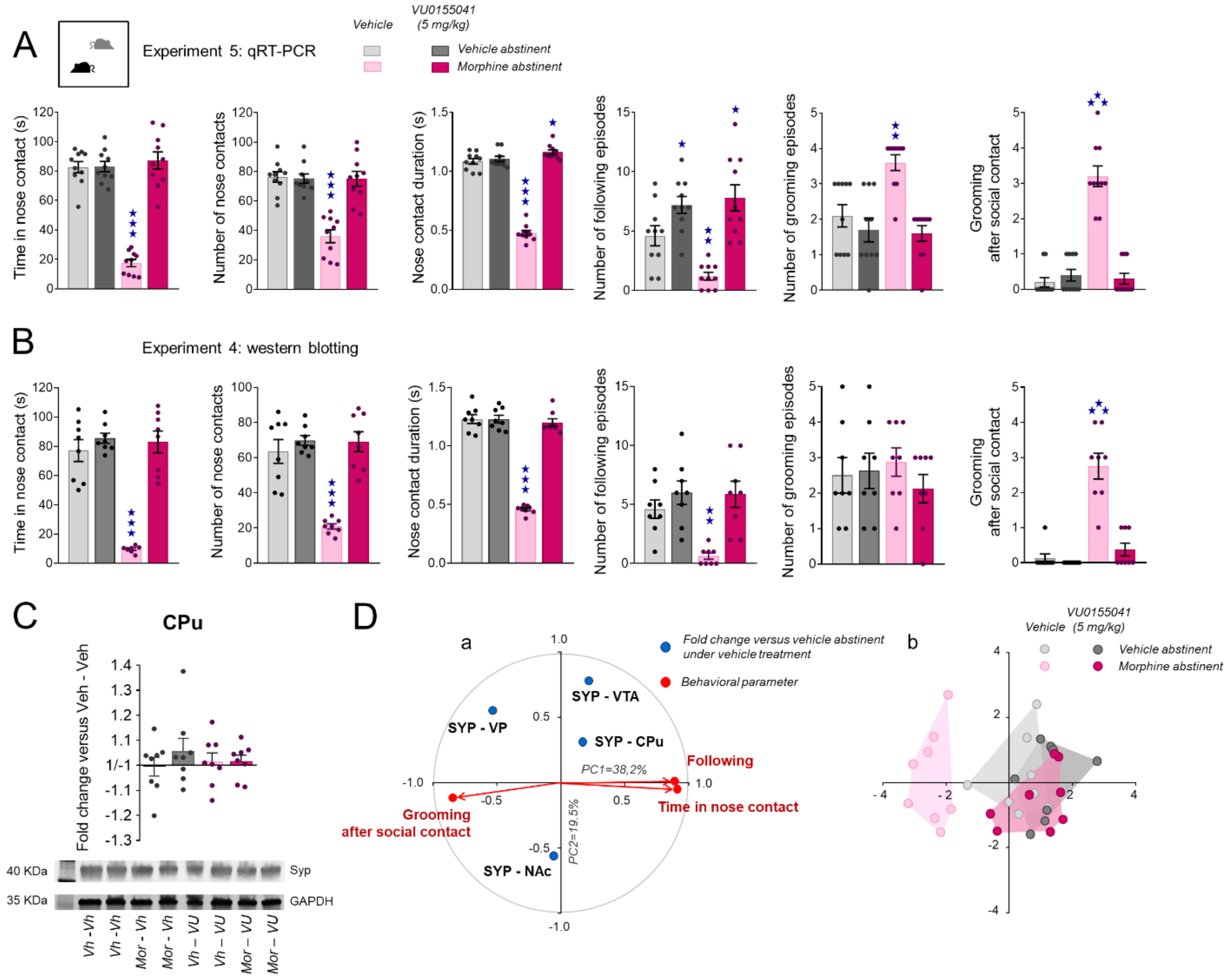


**Figure S3. Effects of chronic VU0155041 treatment of social behavior and protein expression in morphine abstinent mice.**

**(A)** In experiment 4 (see Figure S1), chronic VU0155041 administration (5 mg/kg, IP, once daily) in morphine abstinent mice restored the time spent in nose contacts (abstinence: $F_{1,35}$=55.4, p<0.0001; treatment: $F_{1,35}$=72.7, p<0.0001; abstinence x treatment: $F_{1,35}$=71.0, p<0.0001), their number (abstinence: $F_{1,35}$=23.9, p<0.0001; treatment: $F_{1,35}$=21.2, p<0.0001; abstinence x treatment: $F_{1,35}$=23.4, p<0.0001) and duration (abstinence: $F_{1,35}$=194.2, p<0.0001; treatment: $F_{1,35}$=314.3, p<0.0001; abstinence x treatment: $F_{1,35}$=280.6, p<0.0001) as well as the number of following episodes (abstinence: $F_{1,35}$=3.2, NS; treatment: $F_{1,35}$=32.4, p<0.0001; abstinence x treatment: $F_{1,35}$=6.3, p<0.05) to vehicle abstinence levels, reduced grooming (abstinence: $F_{1,35}$=5.7, p<0.05; treatment: $F_{1,35}$=17.1, p<0.001; abstinence x treatment: $F_{1,35}=7.5$, $p<0.01$) and suppressed grooming after social contact (abstinence: $F_{1,35}=51.8$, $p<0.0001$; treatment: $F_{1,35}=44.7$, $p<0.0001$; abstinence x treatment: $F_{1,35}=59.4$, $p<0.0001$). **(B)** In experiment 5 (see Figure S1), chronic VU0155041 administration (5 mg/kg, IP, once daily) in morphine abstinent mice restored the time spent in nose contacts (abstinence: $F_{1,28}=39.7$, $p<0.0001$; treatment: $F_{1,28}=54.3$, $p<0.0001$; abstinence x treatment: $F_{1,35}=34.4$, $p<0.0001$), their number (abstinence: $F_{1,28}=21.4$, $p<0.0001$; treatment: $F_{1,28}=33.7$, $p<0.0001$; abstinence x treatment: $F_{1,28}=20.0$, $p<0.001$) and duration (abstinence: $F_{1,28}=172.2$, $p<0.0001$; treatment: $F_{1,28}=147.6$, $p<0.0001$; abstinence x treatment: $F_{1,28}=148.1$, $p<0.0001$) as well as the number of following episodes (abstinence: $F_{1,28}=5.9$, $p<0.05$; treatment: $F_{1,28}=15.1$, $p<0.001$; abstinence x treatment: $F_{1,28}=5.2$, $p<0.05$) to vehicle abstinence levels and did not modify grooming globally but suppressed grooming after social contact (abstinence: $F_{1,28}=49.2$, $p<0.0001$; treatment: $F_{1,28}=34.1$, $p<0.0001$; abstinence x treatment: $F_{1,28}=27.7$, $p<0.0001$). **(C)** In experiment 5, levels of SYP in the CPu were not modified by morphine abstinence or chronic VU0155041 treatment (gels in Figure S4 and S5). **(D)** In this same experiment, PCA analysis indicates that SYP expression in the VP was negatively correlated with prosocial parameters in the direct social test (panel a), indicating that individuals with low SYP levels in the VP were more prone to interact with their congeners (panel b). Results are shown as scatter plots and mean ± sem. Solid stars: abstinence x treatment interaction, comparison to the vehicle abstinent group (two-way ANOVA followed by Newman-Keules post-hoc test). One symbol: $p<0.05$, two symbols: $p<0.01$, three symbols: $p<0.001$. NAc: nucleus accumbens, CPu: caudate putamen, PC: principal component, SYP: synapstophysin; Veh: vehicle; VP: ventral pallidum; VU: VU0155041.

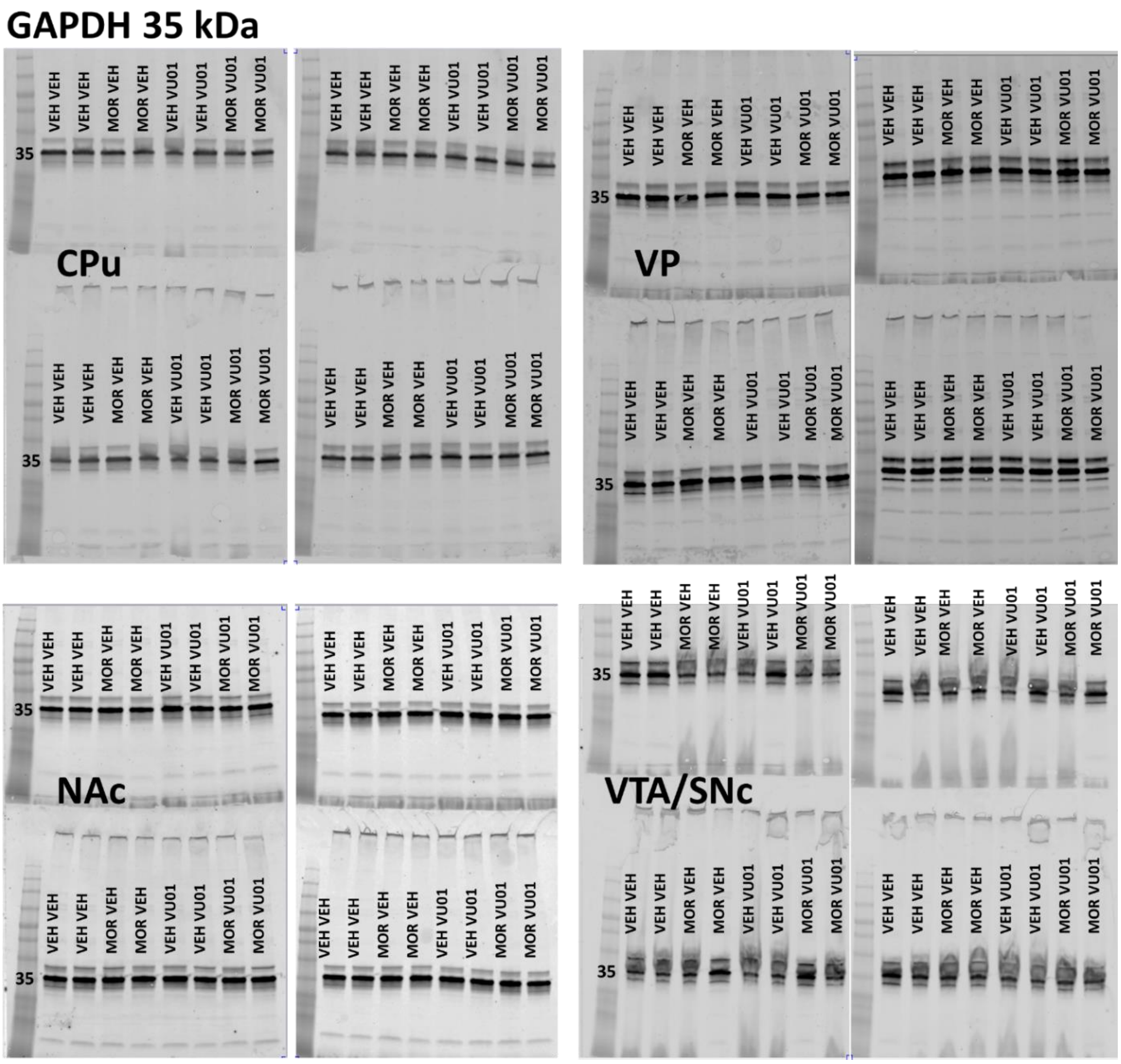


**Figure S4. Gels from western blot experiments:** GAPDH (middle band, 35 kDa) in vehicle-vehicle (VEH-VEH), morphine-vehicle (MOR-VEH), vehicle-VU0155041 (VEH-VU01) and morphine-VU0155041 (MOR-VU01) abstinent mice.

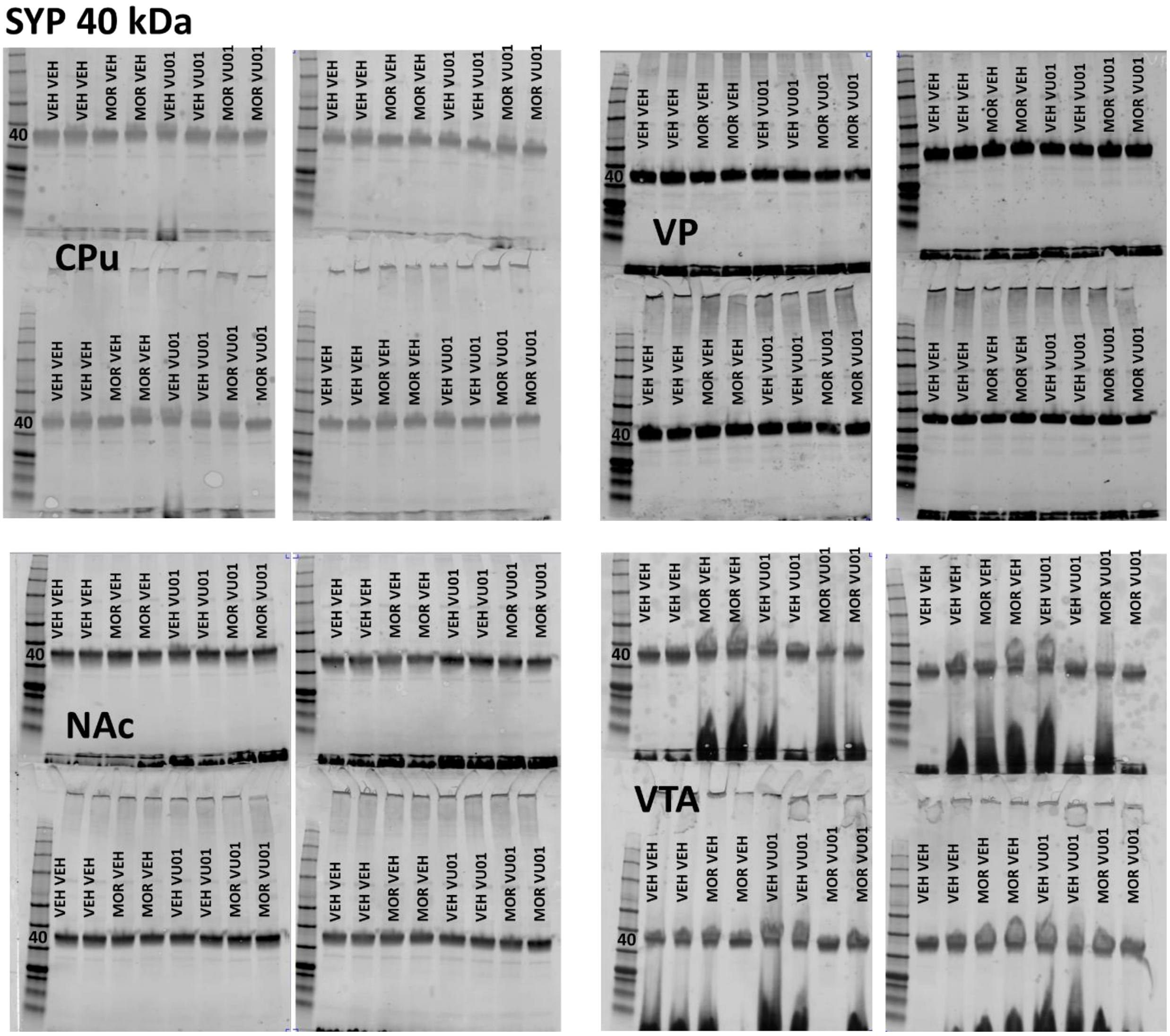


**Figure S5. Gels from western blot experiments:** SYP (middle band, 40 kDa) in vehicle-vehicle (VEH-VEH), morphine-vehicle (MOR-VEH), vehicle-VU0155041 (VEH-VU01) and morphine-VU0155041 (MOR-VU01) abstinent mice.

### Legends to Supplementary Tables

**Table S1. List of primers used for qRT-PCR.**

**Table S2. Transcription levels of a set of 76 genes in the CPu, NAc, BNST and CeA in morphine and cocaine versus vehicle abstinent mice.**

Data are expressed as fold change versus the vehicle abstinent group (median ± SEM). Student’s t-tests were performed on transformed data (see Material and Methods) to determine whether fold changes differed from 0 (no regulation:

corresponds to ± 1.00 in table). Significant regulations of gene expression are highlighted in bold and filled in red or in blue for significant down-regulation. Marker genes of MSNs are quoted in blue; genes of the HTT-related network are underlined. n.d.: not detected.

**Table S3. Transcription levels of a set of 33 genes in the CPu, NAc and CeA in morphine or vehicle abstinent mice treated chronically with VU0155041 or vehicle.**

Data are expressed as fold change versus the vehicle - vehicle abstinent group (median ± SEM). Student's t-tests were performed on transformed data (see Material and Methods) to determine whether fold changes differed from 0 (no regulation: corresponds to ± 1.00 in table). Significant regulations of gene expression are highlighted in bold and filled in red or in blue for significant down-regulation. Marker genes of MSNs are quoted in blue.
